# TRAIL-mediated PMN-MDSC depletion prevents CD4^+^ T cell loss in HIV-infected humanized mice

**DOI:** 10.1101/2025.01.22.634356

**Authors:** Shawn Abeynaike, Kayla Frank, Angel Gandarilla, Silke Paust

**Author notes:** Correspondence: Silke Paust, The Jackson Laboratory for Genomic Medicine 10 Discovery Drive, Farmington, CT 06032.

## Abstract

Myeloid-derived suppressor cells (MDSCs) are critical regulators of HIV immunity, but the mechanisms behind their expansion and value as a therapeutic target remain unclear. We investigated MDSC function in HIV using a humanized mouse model. Acute HIV-1 infection triggered the systemic expansion of polymorphonuclear MDSCs (PMN-MDSCs), with their frequency correlating positively with plasma viral load. Notably, PMN-MDSC expansion increased further when natural killer (NK) cells and cytotoxic T lymphocytes (CTLs) were depleted, suggesting these immune cells restrain MDSC proliferation. We identified Tumor necrosis factor related apoptosis-inducing ligand (TRAIL)-induced apoptosis as a regulatory mechanism, as MDSCs expressed high levels of TRAIL receptor 2 (R2), while NK cells and CTLs upregulated TRAIL expression during HIV infection. Treating HIV-1-infected mice with a TRAIL-R2 agonist significantly reduced MDSC levels to those of uninfected mice and significantly improved NK cell and CTL immune function. Crucially, in viremic humanized mice, the treatment prevented CD4+ T cell loss, a hallmark of HIV-1 pathogenesis and acquired immune deficiency syndrome (AIDS) progression. These findings reveal the therapeutic potential of targeting MDSC apoptosis to mitigate CD4+ T cell depletion and enhance HIV-specific immunity, offering a new clinical target to prevent AIDS and insights into MDSC– NK/CTL cross-regulation during HIV-1 infection.

## Introduction

Over 38.4 million people currently live with human immunodeficiency virus (HIV)-1, and 650,00 HIV-1-related deaths were reported in 2021 ^1^. Untreated HIV-1 infection leads to the progressive depletion of CD4^+^ T cells, the virus’ primary cellular target ^2^. CD4^+^ T cells are critical facilitators of both the humoral and cellular immune response ^3,4^, and their loss leads to acquired immunodeficiency syndrome (AIDS), increasing susceptibility to opportunistic infections and certain cancers ^5^. Although combination antiretroviral therapy (cART) suppresses HIV-1 replication and halts the depletion of CD4^+^ T cells ^6^, it does not eliminate the virus as it persists in long-lived CD4^+^ T cells; thus, people living with HIV-1 (PLHIV) require lifelong treatment with daily oral medications or long-acting injectables ^7^. In addition, some PLHIV are “immunological non-responders” (INRs) to cART and may experience incomplete immune reconstitution and consequential immune impairment ^8^. Even when cART treatment is deemed successful, chronic inflammation and immune dysfunction can persist ^8^, increasing the incidence of co-morbidities ^9^ and co-infections for PLHIV ^10,11^. Teasing apart the roles of innate and adaptive immune cells in this dysregulated state will be essential for developing treatments that can nullify the immune effects of chronic HIV-1 infection.

Cytotoxic lymphocytes such as CD8^+^ T cells (CTLs) and natural killer (NK) cells kill HIV-1-infected cells ^12,13^ by releasing apoptosis-inducing granules (perforin and granzymes), ligating the First Apoptosis Signal (FAS)-ligand or death receptors with FAS or tumor necrosis factor-related apoptosis-inducing ligand (TRAIL), respectively ^12,14^. In addition, NK cells and CTLs release antiviral cytokines such as IFNγ and TNFα ^12^. While many HIV-1-infected cells are cleared by CTLs and NK cells, progression to chronic HIV-1 infection is inevitable and causes generalized, sustained immune activation and subsequent exhaustion, including functional defects in both CTLs ^15^ and NK cells ^12^. One cause of this immune dysfunction is the expansion of myeloid-derived suppressor cells (MDSCs) that begins early in acute infection and persists into chronic infection ^16–26^. Human MDSCs ^27–35^ can be divided into monocytic (M-MDSCs; CD56^-^CD3^-^CD11b^+^CD14^+^HLA-DR^−/lo^CD15^−^) and polymorphonuclear (PMN-MDSCs (CD56^-^CD3^-^CD11b^+^CD33^+^CD14^-^CD15^+^); also known as granulocytic MDSCs), respectively ^36,37^. Initially observed in cancer patients, expansion of MDSCs also occurs in autoimmune disorders ^38–47^, obesity ^48–52^, and pregnancy ^53–63^, as well as during infection by bacteria ^64–68^ or fungi ^69–71^ and in chronic infection with viruses such as hepatitis C virus (HCV) ^72–76^ and HIV-1 ^17–24,26^.

Given the immunosuppressive effects of MDSCs, it is not surprising that their expansion correlates with HIV-1 infection. This expansion is attributed directly to HIV-1 proteins Tat ^21,22^, gp120 ^21^, and Nef ^77^ and indirectly to residual immune activation denoted by elevated levels of the pro-inflammatory cytokine IL-6 and chemokines (CXCL10, CCL4, CXCL8) ^16^. Notably, MDSCs interfere with CD4 T cell reconstitution, possibly by inhibiting the expansion of early hematopoietic progenitors ^17,22,23^. However, PMN-MDSC and CD4^+^CD25^+^FoxP3^+^ regulatory T cell (Treg) levels are positively correlated in chronic HIV-1 infection ^23^, and co-culture of M-MDSCs with CD4^+^ T cells induces Treg differentiation and expansion ^21^. Furthermore, MDSCs from PLHIV suppress CTL cell proliferation ^23,24,26^, and effector functions ^26^. Direct contact with T cells was required for MDSCs from PLHIV to exert their immunosuppressive function ^19,22^.

What is surprising is that MDSC expansion is positively correlated with HIV-1 viral load throughout infection. In combination antiretroviral therapy (cART)-naïve HIV-1-infected individuals, PMN-MDSC frequency is significantly higher than in uninfected individuals, and the frequency of MDSCs is higher in those individuals with progressive viral load (HIV progressors) compared to individuals in which the viral load remains below < 50,000 copies/mL (HIV controllers) ^17,78^. Also, in HIV-infected treatment-naïve persons, PMN-MDSC expansion positively correlated with viral load and negatively correlated with CD4^+^ T cell count. Further, while PMN-MDSC frequency declined with ART, it remains elevated compared to HIV uninfected controls ^17,19,20,23,24,26^. Similarly, studies in other cohorts found elevated levels of M-MDSCs in PLHIV ^16,21,22^. Expansion of MDSCs has also been observed in SIV-infected rhesus macaques ^25,79^. MDSC levels were low before SIV infection then increased during acute infection and persisted during cART ^25^. Interestingly, after cART interruption, the level of MDSCs—particularly PMN-MDSCs (called granulocytic MDSCs in ^25^)— increased substantially and significantly suppressed the proliferation of CD4^+^ and CD8^+^ T cells in response to polyclonal or SIV-specific stimulation *in vitro*. Furthermore, MDSC frequency correlated with the level of circulating inflammatory cytokines, and upon cART interruption, levels of both MDSCs and inflammatory cytokines were more pronounced than in the acute infection. These findings suggest that MDSCs, especially during the post-cART inflammatory cytokine surge, limit immune responses to HIV/SIV infection. This dampening of the antiviral response is a problem as it will undermine the effectiveness of curative strategies ^25^. Although expansion of MDSCs in HIV-1 infection is well supported, findings from several studies have produced somewhat inconsistent results with respect to which MDSC population expands most, perhaps due to differences in infection stage of patients in the study cohorts or factors related to their analyses.

Despite the observations of MDSC expansion during pregnancy, cancer, and infection, the mechanisms regulating MDSC levels and their functional impact are not fully understood. Several reports have suggested involvement of the apoptosis-inducing TRAIL pathway. For example, the level of soluble TRAIL in plasma has been found to be inversely correlated with PMN-MDSC frequency in cART-naïve primary HIV-1-infected patients ^20^, and TRAIL levels were elevated in acute and chronically HIV-1-infected patients compared to healthy donors ^20^. Notably, recombinant TRAIL has been shown to induce apoptosis of PMN-MDSCs isolated from primary HIV-1-infected patients *in vitro* ^20^. Although TRAIL-mediated selective depletion of MDSCs expressing TRAIL death receptors has been demonstrated to have beneficial effects in preclinical models ^80^ and clinical trials for cancer therapy ^81^, the effects and therapeutic value of depleting MDSCs during HIV-1 infection have not yet been investigated.

Using a recently developed humanized immune system mouse model that supports robust human immune cell reconstitution and exhibits antiviral NK cell responses during sustained HIV-1 infection ^82^, we found that HIV-1 infection induces expansion of PMN-MDSCs, consistent with reports in PLHIV and SIV-infected macaques ^17,25^. Depleting NK cells from HIV-1-infected humanized mice led to further PMN-MDSC expansion, suggesting that NK cells inhibit MDSC expansion upon HIV infection. We also discovered increased TRAIL expression on NK cells and T cells upon HIV-1 infection, concomitant with expression of the apoptosis-activating TRAIL receptor 2 (R2) on MDSCs. Building on this finding, we demonstrated that treatment of these HIV-1-infected mice with a TRAIL-R2 agonistic antibody significantly reduced MDSC frequency, correlating with improved degranulation and TNFα production by CTLs and enhanced activation and degranulation of NK cells. A hallmark of HIV-1 infection and pathogenesis is the gradual loss of CD4^+^ T cells with progressive impairment of immunity that ultimately leads to AIDS and death by opportunistic infections or certain cancers. Strikingly, treatment of HIV-1-infected mice with the TRAIL-R2 agonistic antibody prevented such HIV-1-elicited CD4^+^ T cell depletion. These findings holds great promise for the management of human HIV-1 infection, as an existing FDA-approved drug used to deplete MDSCs in cancer settings ^83^ could potentially be repurposed as a therapeutic agent to treat PLHIV.

## Results

### HIV-1 infection induces PMN-MDS expansion in Hu-NSG-Tg(IL-15) mice

We examined HIV-infection induced MDSC expansion in the recently developed Hu-NSG-Tg(IL-15) mouse model, which successfully reconstitutes human NK cells and supports sustained HIV-1 infection ^82^. We used a previously described cytometric gating strategy to identify and quantify human MDSCs in humanized mice ^84^. At 8 weeks post-infection of Hu-NSG-Tg(IL-15) mice with HIV-1 Q23.17 (clade A), we observed an expansion of PMN-MDSCs (CD56^-^CD3^-^CD11b^+^CD33^+^CD14^-^CD15^+^) compared to donor-matched control mice (Figure 1A). This increase in PMN-MDSCs was observed in the peripheral blood, spleen, and liver of HIV-1-infected Hu-NSG-Tg(IL-15) mice (Figure 1A) and was consistent across four different Hu-NSG-Tg(IL-15) cohorts. In previous studies of several human cohorts and SIV-infected rhesus macaques, PMN-MDSC expansion correlated with plasma viral load ^17,25^. Therefore, we compared the percentage of PMN-MDSC against the plasma viral load of HIV-1-infected mice at week eight post infection. We found a significant positive correlation between the plasma viral RNA load and the percentage of PMN-MDSCs in the peripheral blood (r=0.063, p=0.021) and liver (r=0.061, p=0.026) but not in the spleen (Figure 1B).

**Figure 1:**
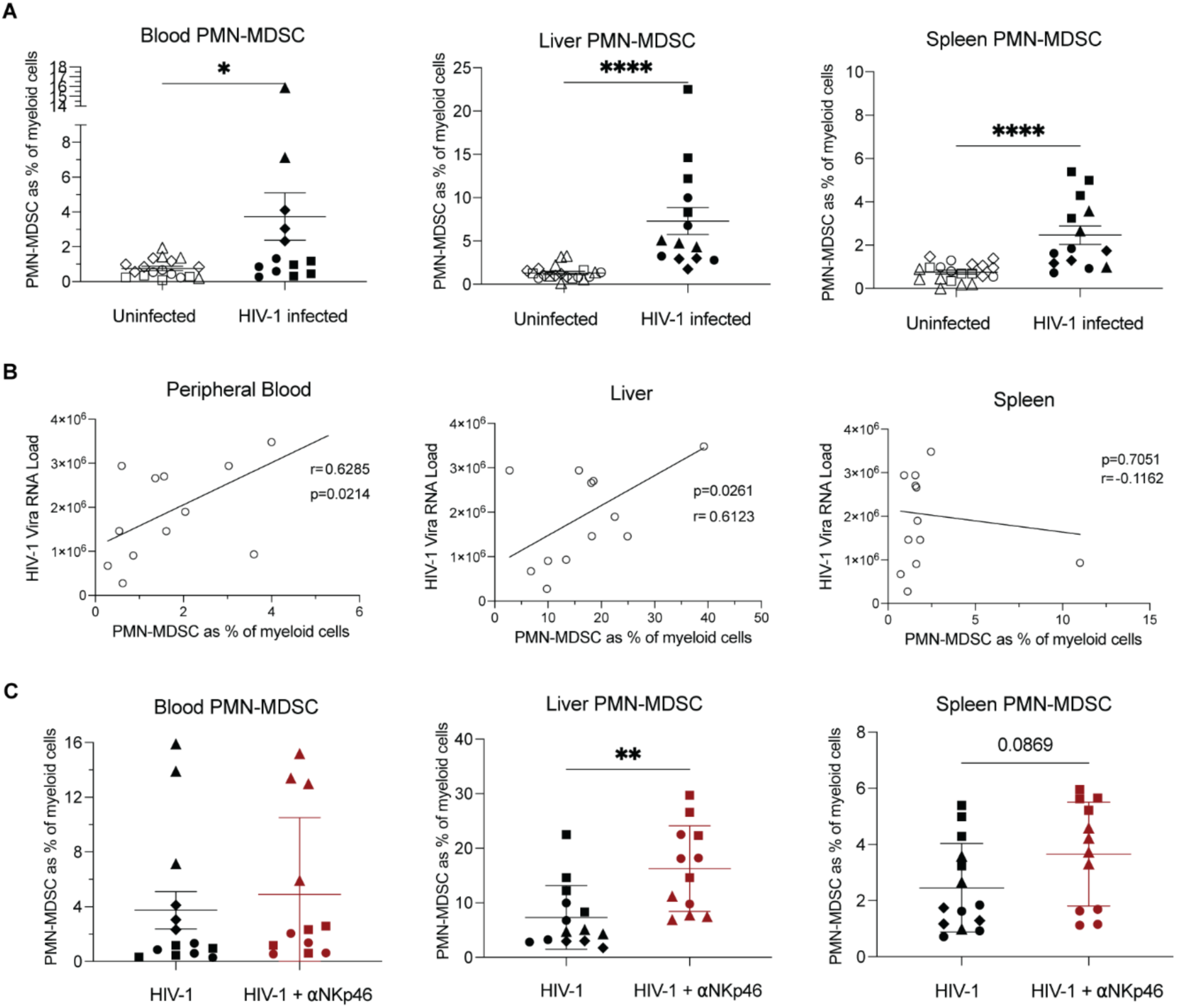
PMN-MDSCs expand during HIV-1 infection. A) Frequency of PMN-MDSC (CD3^-^CD56^-^CD11b^+^HLADR^low/-^CD33^+^CD14^-^CD15^+^) in blood, liver, and spleen of Hu-NSG-Tg(IL-15) mice 8 weeks post intraperitoneal infection with Q23.17 (10^5^ IU). Data is presented of individual mice from four human donor-matched cohorts represented by matching symbols (n=3-4 mice per donor, per experimental group). Statistical significance was calculated using an unpaired t-test *P < 0.05, **P < 0.01, **P < 0.005 and ****P < 0.0001. B) The percentage and number of PMN-MDSC from the blood, liver and spleen 8 weeks post-infection positively correlate with viral RNA load in the plasma of HIV-1 infected Hu-NSG-Tg(IL-15) mice. Correlation of the percentage of PMN-MDSC of total myeloid cells (CD3^-^CD56^-^CD11b^+^) as determined by flow cytometry and viral RNA load of HIV-infected hu-IL-15Tg NSG mice at 8 weeks post-infection. C) Frequency of PMN-MDSC in blood, liver and spleen of HIV-1 infected mice with or without depletion of NK cells by weekly intraperitoneal injection of Anti-Nkp46 (9E2). Data is presented from individual mice from three human donor-matched cohorts represented by matching symbols (n=4 per donor, per experimental group). Statistical significance was calculated using an unpaired t-test *P < 0.05, **P < 0.01, **P < 0.005, and ****P < 0.0001.

NK cells can kill MDSCs in humanized mouse models of solid tumor ^85^. Thus, we utilized an antibody specific to Nkp46 (clone 9E2) to deplete NK cells in HIV-1-infected Hu-NSG(IL-15Tg) mice to determine whether NK cells impact HIV-induced MDSC expansion, and, if so, to quantify such an impact. Of note, because the NKp46-specific antibody clone 9E2 recognizes the domain 1-NKp46 splice variants and does not recognize the domain 2 extracellular domain ^86^, it results in a statistically significant reduction of NK cells (CD56^+^CD3^-^) but not complete depletion (Supplemental Figure 1A). Nevertheless, NK cell reduction further increased PMN-MDSC frequency in the liver of HIV-1-infected mice compared to HIV-1-infected donor-matched control mice (Figure 1C). Next, we depleted CD8a-expressing cells (CTL and CD8a^+^ NK) in Hu-NSG(IL-15Tg) mice and discovered an increased PMN-MDSC frequency in the liver but not spleen or peripheral blood (Supplemental Figure 1B). The targeting of CD8a-positive cells depleted CTLs as expected, as well as a subset of NK cells (Supplemental Figure 1 and ^87^). Our results suggest that NK cells and CTLs inhibit the expansion of MDSCs upon HIV-1 infection.

### Inhibition of PMN-MDSC expansion by NK cells and CTLs via the TRAIL pathway

In PLHIV, PMN-MDSC frequency inversely correlates with plasma TRAIL levels, and recombinant TRAIL can induce the apoptosis of PMN-MDSCs isolated from PLHIV ^20^. Therefore, we experimentally evaluated if the TRAIL pathway inhibits MDSC expansion in acute HIV-1 infection. Indeed, we observed high levels of TRAIL-R2 expression on PMN-MDSCs and M-MDSCs in the peripheral blood, liver, and spleen of HIV-1-infected Hu-NSG-Tg(IL-15) mice (Figure 2A, B). We also observed the expression of TRAIL-R1 on both PMN-MDSCs and M-MDSCs (Figure 2A, B), albeit to a lesser extent.

**Figure 2:**
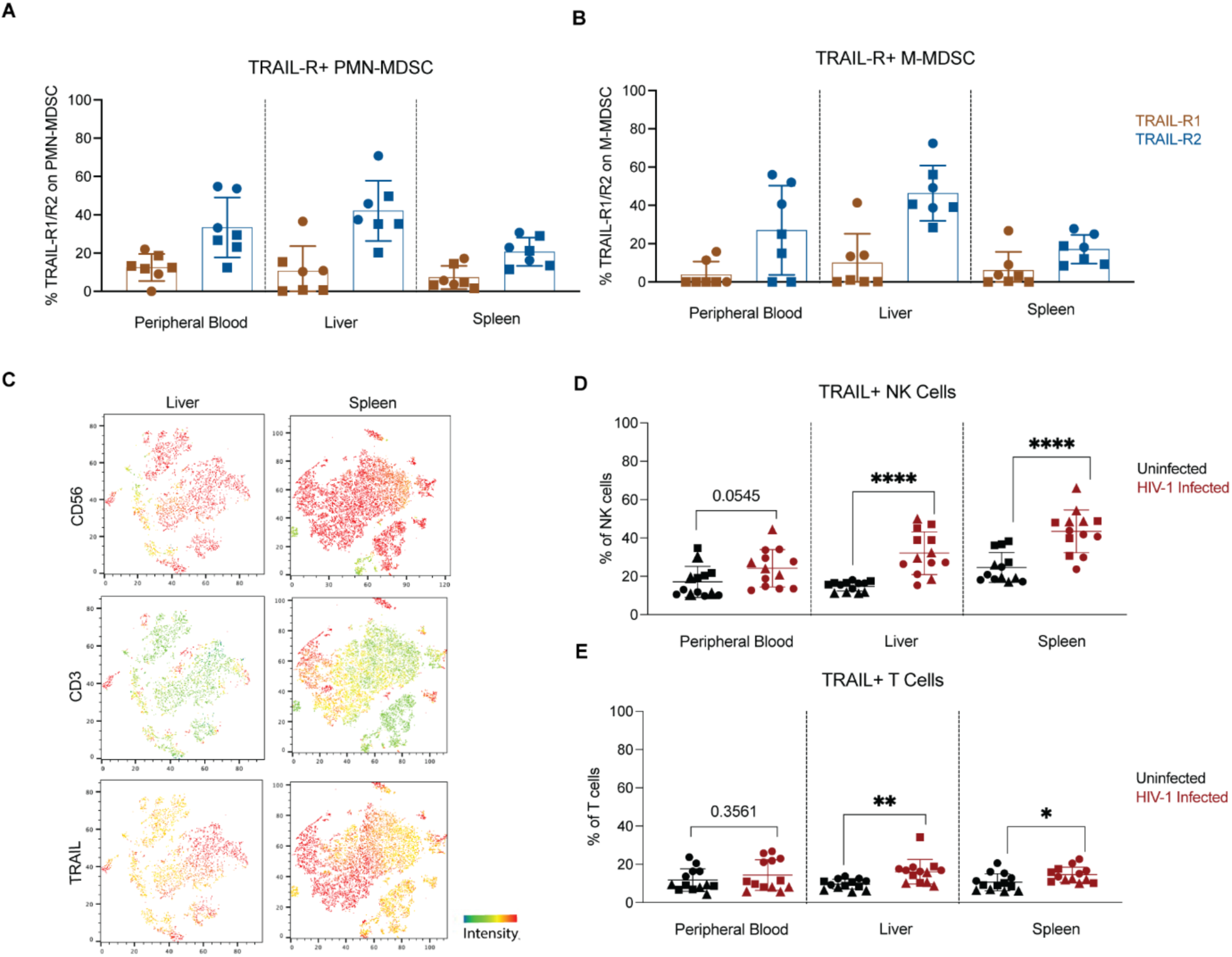
Expression of TRAIL on NK cells and T Cells and TRAIL receptors on PMN-MDSCs during HIV-1 infection. A-B) Percentage of TRAIL-R1 and TRAIL-R2 expression on PMN-MDSCs and M-MDSCs in HIV-1 infected mice (2 donors, n=3-4 per group). C) t-SNE analysis of mCD45^-^hCD45^+^TRAIL^+^ cells in HIV-1 infected mice (1 donor, n=4 mice per donor) overlayed with expression heatmaps of CD56, CD3 and TRAIL. D) Percentage of TRAIL expressing NK cells in blood, liver and spleens of Hu-NSG-Tg(IL-15) mice 8 weeks post intraperitoneal infection with Q23.17 (10^5^ IU) and uninfected human donor-matched controls. (3 donors, n=3-4 mice per donor, per experimental group). E) Frequency of TRAIL expressing T cells in blood, liver and spleens of Hu-NSG-Tg(IL-15) mice 8 weeks post intraperitoneal infection with Q23.17 (10^5^ IU) and uninfected human donor-matched controls. (3 donors, n=3-4 mice per donor, per experimental group). Statistical significance was calculated using unpaired t-tests. *P < 0.05, **P < 0.01, **P < 0.005 and ****P < 0.0001.

To dissect the composition of TRAIL^+^ human immune cells (hCD45^+^mCD45^-^) in Hu-NSG-Tg(IL-15) mice, we performed t-distributed stochastic neighbor embedding (t-SNE) analysis. The majority of TRAIL-producing cells were NK cells (CD56^+^CD3^-^) (Figure 2C), which demonstrated a substantial increase in surface expression of TRAIL (Figure 2D), both in the livers and spleens of Hu-NSG-Tg(IL-15) mice at eight weeks post HIV-1 infection, compared to uninfected human donor-matched control mice. A similar trend was seen for NK cells in the peripheral blood but did not reach statistical significance (p=0.0545). Likewise, a modest but statistically significant increase in TRAIL expression was seen on T cells (CD3^+^CD56^-^) in the liver and spleen, but not peripheral blood, of HIV-1-infected Hu-NSG-Tg(IL-15) mice compared to uninfected donor-matched controls (Figure 2E). Thus, during acute HIV-1 infection, TRAIL-expressing NK cells and T cells inhibit the expansion of MDSCs; however, as immune exhaustion and dysfunction ensue, this mechanism is not sufficient to fully inhibit MDSC expansion.

### Treatment with TRAIL-R2 agonistic antibody leads to the depletion of MDSCs and mitigates CD4^+^ T cell loss

Given the high expression of TRAIL-R2 on MDSCs in HIV-1-infected Hu-NSG-Tg(IL-15) mice, we tested if the TRAIL pathway could be exploited to deplete MDSCs *in vivo*. We treated HIV-1-infected Hu-NSG-Tg(IL15) mice with a TRAIL-R2 agonistic antibody (MAB6313) that has been previously shown to selectively induce apoptosis of human MDSCs expressing TRAIL-R2 *in vivo* ^81^. We observed a decrease in PMN-MDSC-frequency in the livers but not the spleens of HIV-1-infected mice treated with TRAIL-R2 agonistic antibody (Figure 3A, B). The percentages and total numbers of M-MDSCs were also significantly reduced in the spleens of HIV-1-infected Hu-NSG-Tg(IL15) mice (Figure 3C), with a similar trend seen in the livers (Figure 3D). Importantly, while weekly intraperitoneal injections with TRAIL-R2 agonistic antibody did not result in significant changes in HIV-1 plasma viral RNA load (Figure 3E, F), HIV-1 infected humanized mice treated with TRAIL-R2 agonistic antibody therapy were resistant to HIV-1-induced CD4^+^ T cell depletion (Figure 3G, H) compared to cohort matched HIV-1-infected untreated control mice, which consistently displayed significant depletion of CD4^+^ T cells. This experiment provides proof of concept that TRAIL-mediated depletion of MDSCs using TRAIL-R2 agonistic antibodies has the potential to protect CD4^+^ T cells from HIV-1-induced depletion in humans.

**Figure 3:**
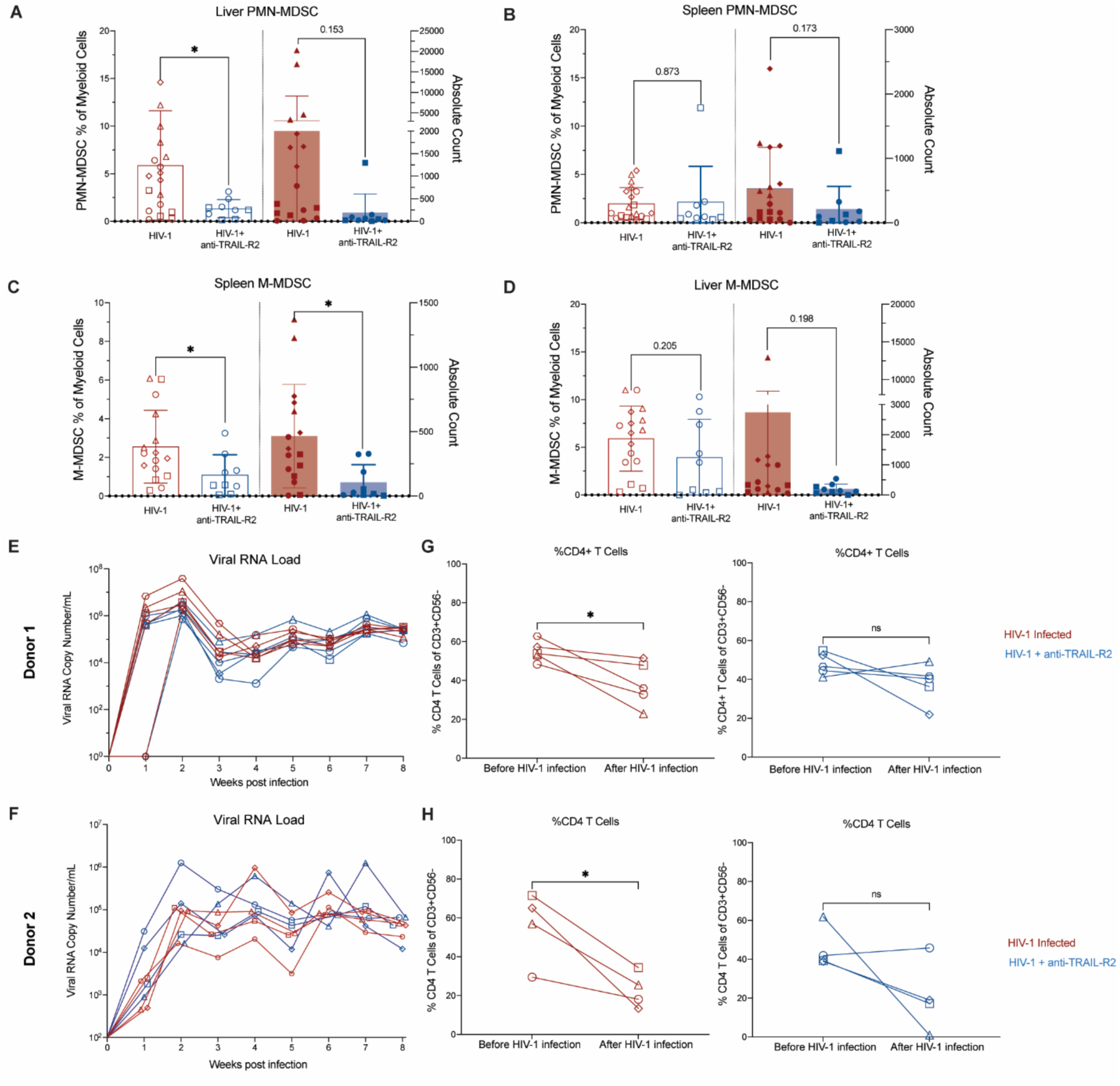
Treatment with anti-TRAIL-R2 antibodies leads to depletion of MDSCs and improved CD4+ T cell reconstitution. **A, B)** Percentage and Total numbers of PMN-MDSCs in the livers and spleens of Hu-NSG-Tg(IL-15) mice 8 weeks post HIV-1 infection, with or without treatment with anti-TRAIL-R2 agonistic antibodies. Data is presented from individual mice from 2-4 human donor-matched cohorts represented by matching symbols (n=3-5 per donor, per experimental group). **C, D)** Percentage and Total numbers of M-MDSCs in the livers and spleens of Hu-NSG-Tg(IL-15) mice 8 weeks post HIV-1 infection, with or without treatment with anti-TRAIL-R2 agonistic antibodies. Data is presented from individual mice from 2-4 human donor-matched cohorts represented by matching symbols (n=3-5 per donor, per experimental group). Statistical significance was calculated using unpaired t-tests. *P < 0.05. **E, F)** Plasma viral RNA load of Hu-NSG-Tg(IL-15) mice infected with Q23.17 and treated with anti-TRAIL-R2 or untreated donor-matched control mice. Data from each individual mouse is depicted by a different symbol (2 donors, n=4-5 per experimental group, per donor). **G, H)** Percentage of CD4+ T cells of total T cells in the peripheral blood before and after HIV-1 infection in anti-TRAIL-R2 treated and untreated donor matched Hu-NSG-Tg(IL-15) mice. Data from each individual mouse is depicted by a different symbol (2 donors, n=4-5 per experimental group, per donor). Statistical significance was calculated using paired t-tests. *P < 0.05.

### Depletion of MDSCs with TRAIL-R2 agonistic antibody leads to NK cell activation and degranulation and CD8+ T cell increased TNFα expression and degranulation

To understand how TRAIL-mediated MDSC depletion modulates cytotoxic lymphocyte functions in HIV-1-infected humanized mice, we performed multispectral flow cytometry focusing on functional and phenotypic markers of splenic NK cells and T cells eight weeks post-infection. We examined the expression of activating receptors (NKG2D and CD69), inhibitory receptors (NKG2A and PD-1), cytotoxic granules (perforin and granzyme B), anti-viral cytokines (IFNγ and TNFα), the Fc receptor CD16/FcgRIII, and transcription factors (T-BET and EOMES), as well as markers of tissue residency (CD69 and CXCR6), maturity (CD57), degranulation (CD107a), and proliferation (Ki67). For improved visualization of human NK cells, T cells, and natural killer T (NKT) cells (CD56+ CD3+) in the flow cytometric data, we performed dimensionality reduction by applying t-SNE to immune cells (hCD45^+^mCD45^-^) that also expressed either CD56 and/or CD3. We additionally performed unsupervised clustering using the FlowSOM (self-organizing maps) algorithm, which revealed nine distinguishable cell population, Pops 1–9.

Pop 1 and Pop 2 exhibit markers for CD4^+^ T cells (CD3^+^CD56^-^CD8^-^) and were both higher in TRAIL-R2 agonistic antibody treated HIV-1-infected mice (Pop1-41.7%, Pop2-1.25%) compared to the uninfected and untreated controls (Pop1 40.1%, Pop2 1.02) (Supplemental Figure 2). Pop 1 is high in T-BET, Ki67, IFNg, and CD107a and low in PD-1, suggesting a Th1 phenotype, while Pop 2 was low in T-BET, but high in EOMES and CD69, suggesting a more tissue-resident memory phenotype. This further validates the improved reconstitution of CD4^+^ T cells upon TRAIL-R2 agonistic antibody treatment.

Importantly, upon treatment with TRAIL-R2 agonistic antibody, we saw an expansion of Pop 3, which contains NK cells (CD56^+^CD3^-^) (Figure 4). This population expressed high levels of CD16, Ki67, IFNg, NKG2D, T-BET, perforin, and granzyme B, suggesting this is an activated, proliferating population of NK cells that is capable of cytolysis, cytokine production, and antibody-dependent cellular cytotoxicity. Hierarchical gating and analysis of NK cells showed a significant upregulation of the NK cell-activating receptor NKG2D (Figure 4D) and the degranulation marker CD107a (Figure 4E) upon TRAIL-R2 agonistic antibody treatment. Additionally, while the percentage of NK cells expressing the proliferation marker Ki67 was not significantly higher in HIV-1-infected mice compared to uninfected controls, the percentage of Ki67^+^ NK cells was significantly higher in the TRAIL-R2 agonistic antibody-treated HIV-1-infected, indicating improved proliferative capacity (Figure 4G).

**Figure 4:**
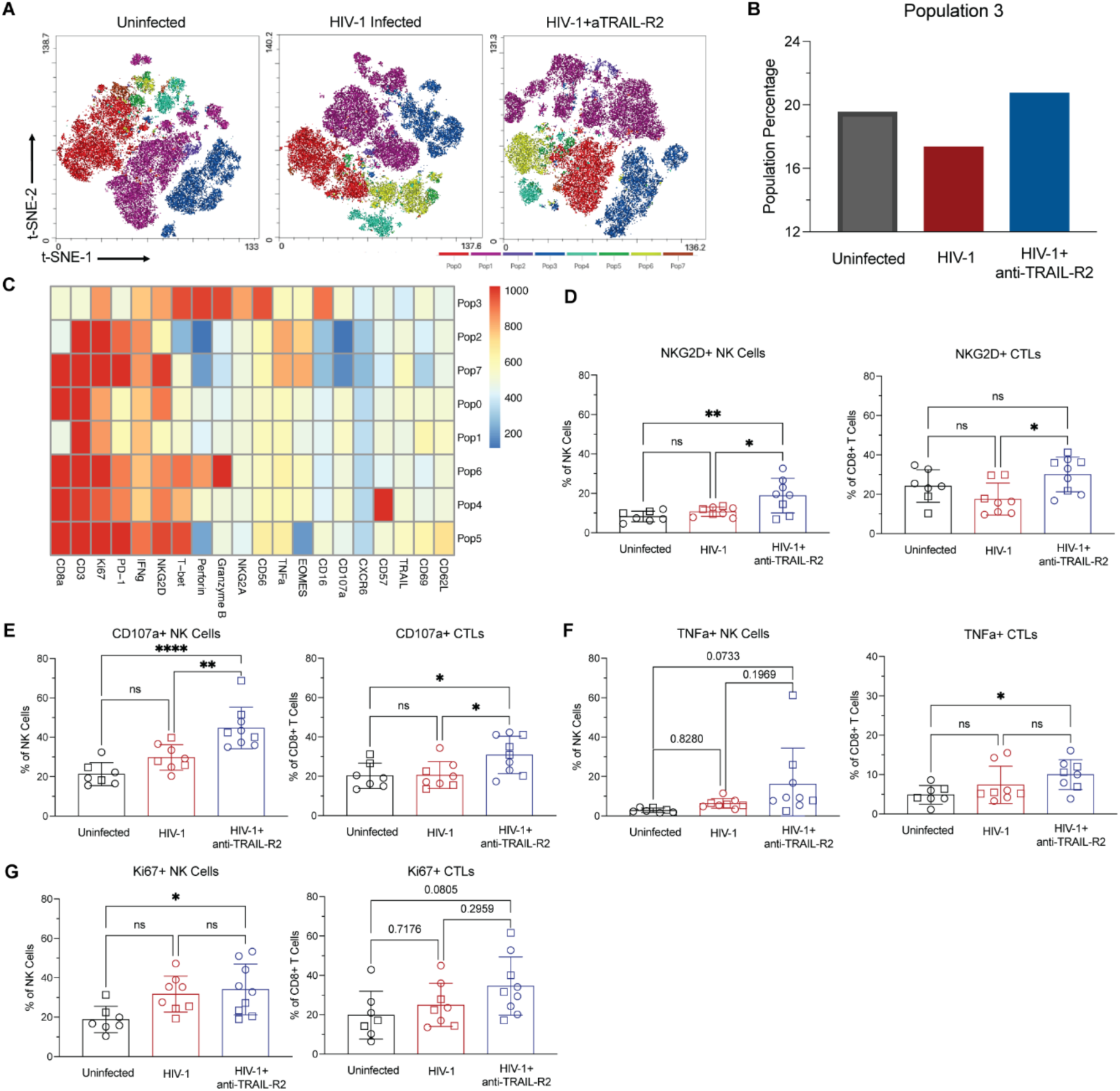
Downstream phenotypic effects on immune cells upon anti-TRAIL-R2 treatment of HIV-1 infected humanized mice. A) t-SNE dimensionality reduction of hCD45+mCD45-cells positive for either CD3 and/or CD56 with overlayed with FlowSOM clusters of splenic immune cells Hu-NSG-Tg(IL-15) mice 8-weeks post HIV-1 infection, with or without treatment with anti-TRAIL-R2 agonistic antibodies and uninfected donor-matched control mice. Cells from individual mice were pooled and downsampled to 50,000 events prior to t-SNE and FlowSOM analysis B) Percentage of cells in Cluster 3 (NK Cells) generated from unsupervised FlowSOM clustering. C) Heatmap displaying phenotypic marker expression for each population clustered using FlowSOM algorithm. D-E) Extracellular expression of NKG2D and CD107a (Lamp-1) on NK cells (CD56^+^CD3^-^) and CTLs (CD56^-^CD3^+^CD8^+^) in Hu-NSG-Tg(IL-15) mice 8-weeks post HIV-1 infection, with or without treatment with anti-TRAIL-R2 agonistic antibodies and uninfected donor-matched controls (2 donors, n=3-5 mice per donor, per experimental group). F-G) Intracellular expression of TNFa and intranucleuar expression of Ki67 on NK cells (CD56^+^CD3^-^) and CTLs (CD56^-^CD3^+^CD8^+^) in Hu-NSG-Tg(IL-15) mice 8-weeks post HIV-1 infection, with or without treatment with anti-TRAIL-R2 agonistic antibodies and uninfected donor-matched controls (2 donors, n=3-5 mice per donor, per experimental group). Statistical significance was calculated using an unpaired t-test *P < 0.05, **P < 0.01, **P < 0.005, and ****P < 0.0001.

Pop 5 contains both NKT cells and CTLs, and the remaining Pops (0, 4, 6, and 7) exhibited CTL phenotypes (Supplemental Figure 2). Hierarchical gating and analysis of CTLs revealed increased expression of CD107a (Figure 4E), indicative of improved degranulation, in spleens of HIV-1-infected mice treated with TRAIL-R2 agonistic antibody compared to both uninfected and HIV-1-infected control mice. Additionally, intracellular TNF*α* expression was higher in CTLs from TRAIL-R2 agonistic antibody -treated HIV-1-infected mice compared to uninfected controls but not untreated HIV-1-infected mice (Figure 4F). Finally, expression of NKG2D on CTLs was higher in TRAIL-R2 agonistic antibody treated mice compared to HIV-1-infected controls (Figure 4D), indicating improved co-stimulation of CTLs.

This analysis highlights several key changes in the splenic immune cell landscape in HIV-1-infected mice that support an antiviral response. Together, our findings demonstrate that depleting MDSCs during HIV-1 infection has the potential to improve NK cell and T cell function.

## Discussion

Acute HIV-1 infection induces inflammatory stimuli that lead to the aberrant expansion and differentiation of monocytes and neutrophils into M-MDSCs and PMN-MDSCs, respectively ^17–24,26^. MDSCs subsequently contribute to immune dysfunction in PLHIV, which is not corrected by cART treatment ^17–24,26^. Here, we investigated the regulation of PMN-MDSC expansion by cytotoxic lymphocytes and explored TRAIL-focused therapeutic approaches to prevent HIV-associated immune dysfunction using the humanized mouse model Hu-NSG-Tg(IL-15). Our study generated several novel and important findings: First, during acute HIV-1 infection in Hu-NSG-Tg(IL-15) mice, the frequency of PMN-MDSCs increased to a level significantly greater than that in uninfected donor-matched control. Depletion of NK cells and CTLs led to further expansion of PMN-MDSCs, identifying cytotoxic cells as regulators of their expansion. Second, TRAIL expression on NK and T cells increased upon HIV-1 infection, and MDSCs expressed high levels of TRAIL-R2. In testing the significance of this, we demonstrated that treatment with a TRAIL-R2 agonist significantly reduced PMN-MDSC frequency. Most importantly, this reduction prevented HIV-1-induced CD4^+^ T cell depletion; enhanced the activation, proliferation, and degranulation of NK cells; and improved TNFα production and degranulation of CTLs. Our findings suggest that the therapeutic depletion of PMN-MDSCs through TRAIL-mediated apoptosis could be a promising therapeutic strategy to mitigate HIV-1-induced immune dysfunction and prevent CD4^+^ T cell depletion, thereby improving immunity in PLHIV.

Although MDSC expansion is an established feature of human HIV-1 infection, it has been challenging to study because it is not exhibited in any previous *in vivo* model of HIV-1 infection. To date, the closest *in vivo* approximation has been MDSC expansion in rhesus macaques in response to related viruses such as SIV ^25,79^. In the present study, we made a significant observation of PMN-MDSC expansion during HIV-1 infection in Hu-NSG-Tg(IL-15) mice. Notably, this expansion was observed in multiple tissues, including the peripheral blood, spleen, and liver, suggesting a systemic impact of HIV-1 infection on MDSC populations. Interestingly, the increase in PMN-MDSC frequency correlated positively with plasma viral load in the peripheral blood and liver, suggesting a potential relationship between MDSC expansion and viral replication. These findings align with previous studies demonstrating PMN-MDSC expansion in human cohorts ^17,19,20,23,24,26^ and SIV-infected macaques (84). We also explored the mechanisms regulating MDSC expansion during HIV-1 infection. Depletion of NK cells further increased PMN-MDSC frequency, indicating a negative regulatory role of these immune cell subsets. Previous studies have demonstrated the capability of MDSCs to suppress NK cells and cytotoxic T lymphocytes (CTLs) in various contexts, such as pregnancy, chronic viral infections, and cancer ^18,24,52,54,59,60,78,88^. Notably, a study of patient-derived tumor samples in xenograft mouse models and has shown that NK cells can effectively eliminate MDSCs in the tumor microenvironment ^85^. Our data suggest that this regulatory mechanism is not restricted to cancer but also applies to acute viral infection. Our findings further support a reciprocal relationship wherein cytotoxic cells also regulate MDSC expansion.

In investigating mechanisms that might underlie the negative regulation of MDSC expansion, we focused on the TRAIL pathway since it has been implicated in the MDSC killing in the contexts of pregnancy and cancer ^81,89^. The expression of TRAIL receptors on PMN-MDSCs and M-MDSCs and an increase in TRAIL expression on NK cells and T cells in HIV infected Hu-NSG-Tg(IL-15) mice suggests a potential role for TRAIL in modulating MDSC expansion. Future studies are needed to determine whether immune exhaustion and dysfunction during HIV infection cause NK cell and T cell-mediated regulation of MDSC expansion to become insufficient. To demonstrate the role of the TRAIL in regulating MDSC expansion during acute infection, we infused HIV-1-infected Hu-NSG-Tg(IL-15) mice with TRAIL-R2 agonistic antibody. Treatment with this antibody resulted in the depletion of both PMN-MDSCs and M-MDSCs in the spleen and liver of HIV-1-infected Hu-NSG-Tg(IL-15) mice. These findings provide compelling evidence that TRAIL-R2 agonistic antibody therapy can successfully deplete MDSCs during HIV-1 infection *in vivo*.

Importantly, the depletion of MDSCs was accompanied by notable changes in the immune cell landscape. A critical indicator of disease progression in HIV-1-infection is the gradual loss of CD4^+^ T cells and immune impairment. T follicular helper (Tfh) cells, a type of Th cell, provide essential support to B cells through direct interactions through CD40–CD40L, cytokine secretion (IL-4, IL-21), and chemoattraction (CXCL13 binding to CXCR5) ^4^. These interactions are crucial for plasma cell differentiation, proliferation, antibody production, class-switch recombination, and germinal center formation. Moreover, CD4^+^ T cells play a vital role in bolstering the primary CD8^+^ T cell response, both increasing cell number and improving functionality ^3^. Therefore, it is noteworthy and clinically relevant that therapeutic TRAIL-R2-mediated PMN-MDSC depletion in HIV-1-infected mice prevented CD4^+^ T cell depletion. This discovery aligns with multiple clinical studies of PLHIV treated with ART where an inverse correlation was found between PMN-MDSC frequency and CD4^+^ T cell count, even after adjusting for age and baseline CD4^+^ T cell count ^17,22,23^. In fact, one publication suggests that PMN-MDSCs may contribute to CD4^+^ T cell depletion by directly inhibiting the expansion of early T cell progenitors^17^. Our investigation identifies TRAIL-mediated killing of MDSCs by cytotoxic T cells as a potential mechanism of MDSC regulation and extends these clinical observations by demonstrating that MDSC depletion enhances CD4^+^ T cell reconstitution *in vivo*.

The NK cell expressed activating receptor NKG2D binds stress ligands MICA/MICB and UL16 binding proteins (ULBPs) 1-6, and plays a crucial role in NK cell-mediated killing of HIV-1^+^ CD4^+^ T cells ^90^. NKG2D also acts as a co-stimulatory receptor on CTLs aiding in the recognition of target cells and enhancing T-cell receptor signaling ^91^. During acute HIV-1 infection, the viral protein Vpr upregulates cell-surface expression of MICA and ULBPs, promoting NKG2D-mediated activation of NK cells ^92,93^. However, in chronic HIV-1 infection, chronic lymphocyte activation causes direct NKG2D downregulation, Nef and Vpu downregulate NKG2D ligands on infected cells, and matrix metalloproteinases cleave NKG2D ligands from the surface of HIV-1-infected CD4^+^ T cells, preventing their killing and resulting in the accumulation of soluble NKG2D ligands that inactivate NKG2D on circulating effector cells ^94,95^. It is therefore noteworthy that we observed upregulation of NKG2D on both NK cells and CTLs upon therapeutic TRAIL-R2-mediated depletion of MDSCs in HIV-1-infected Hu-NSG-Tg(IL-15) mice, suggesting improved activation and co-stimulation of these effector cells respectively. Further, upon therapeutic TRAIL-R2 targeting of HIV-1-infected mice, we observed a significant increase in lysosome-associated membrane protein 1 (LAMP1)/CD107a on the surface of both NK cells and CTLs, reflecting the fusion of lytic granule-containing lysosomes with the plasma membrane ^96,97^ and indicating enhanced degranulation and, therefore, increased cytolytic function. Moreover, NK cells and CTLs produce cytokines such as IFNγ and TNFα that can limit viral replication ^12,98^. TRAIL-R2 agonistic antibody therapy also increased intracellular expression of TNFα on splenic CD8^+^ T cells compared to untreated donor-matched control mice. A similar trend in TNFα expression was observed in NK cells, although it did not reach statistical significance. This is important, since TNFα can inhibit HIV-1 entry, primarily through the downregulation of the HIV CD4 receptor and CCR5 co-receptor ^99^. Further, TNF signaling through its receptor elicits the secretion of factors that suppress HIV-1 replication, such as macrophage inflammatory protein-1α (MIP-1α), MIP-1β, and regulated upon activation, normal T cell expressed and secreted (RANTES) ^100^. Notably, HIV-1 Tat vaccines induce CD8^+^ T cell production of TNFα, which correlates with vaccine-mediated protection ^101,102^, the increase of CD4^+^ T cells and reduced proviral load in clinical trials ^103–105^.

An interesting observation in published clinical data of PLHV cohorts is that while most studies have reported expansion of PMN-MDSCs ^17,19,20,23,24,26^, some studies reported expansion of M-MDSCs ^16,21,22^. The factors underlying this difference remain unknown, but it could be due to variation in HIV-1 strains, host genetic factors, microbiome composition, diet, and/or socioeconomic factors. In this context, humanized mouse models offer a significant experimental advantage as they enable the targeting of MDSCs during HIV-1 infection in a highly controlled setting, whereby genetically identical NSG mice that share identical housing and feed are used to generate multiple cohorts of humanized mice, whereby each cohort is generated using HSCs from a single cord blood unit ^106^. However, humanized mouse models also have limitations, including that T cells in the Hu-NSG-Tg(IL-15) mouse model are not human leukocyte antigen (HLA) restricted. However, even non-HLA-restricted CD8^+^ T cells can respond to HIV-1 peptide pools by producing antiviral cytokines ^107^, and CTL-mediated killing by TRAIL does not require HLA-recognition, but recognition of DRs4 or 5.

Multiple strategies have been pursued to develop a functional or sterilizing cure for HIV-1 infection. Among these, the “shock-and-kill” strategy aims to disrupt HIV-1 viral latency to subsequently eliminate the reservoir of infected cells ^108^. However, this strategy alone has proven inadequate in achieving efficient viral clearance due to the compromised immune response ^108^. Therefore, additional interventions such as therapeutic vaccines, broadly neutralizing antibodies, or infusion of genetically engineered T cells or NK cells will be required to augment the “kill” arm. Encouragingly, therefore, the utilization of TRAIL-R2 agonistic antibodies could serve as a valuable adjunct to reinforce the T cell or NK cell response following therapeutic vaccines, immune checkpoint blockade, or cell infusions in the context of HIV-1 as well. Another approach, termed the “block-and-lock” strategy, aims to suppress HIV-1 replication and induce a state of deep latency, offering the potential for a functional cure ^109^. In this context, adjunctive therapy with a TRAIL-R2 agonistic antibody following induction of this deep latent state may help ameliorate immune dysfunction and restore a more physiologically balanced immune landscape.

The expansion of MDSCs has been identified in mono-infection with HCV ^72,74,88^ or active Mtb ^110,111^. PLHIV are at greater risk of co-infection with hepatitis C virus (HCV) or *Mycobacterium tuberculosis* (Mtb) ^112^ and have a higher risk of reactivation of latent Mtb even with successful antiretroviral therapy ^112,113^, primarily due to the immune dysfunction associated with chronic HIV-1 infection, which includes depletion and suppression of Mtb-specific T cells that may be in part due to MDSC expansion ^11,112,113^. In the context of HCV infection, co-culture experiments have demonstrated the ability of MDSCs to inhibit IFN-γ release from NK cells ^72^ and impair autologous T-cell function ^73^. Furthermore, despite the ability of direct-acting antivirals to cure HCV infection in both HCV mono-infected ^74^ and HCV/HIV-1 co-infected patients ^114^, the expanded population of MDSCs persists and contributes to generalized immune suppression. Depletion of MDSCs through TRAIL-R2 agonistic antibody therapy could serve as a valuable tool to combat these co-infections in PLHIV in conjunction with traditional pharmacological prophylactic and curative treatments. Importantly, targeting of TRAIL-receptors is likely safe for humans, as a clinical trial involving patients with solid tumors demonstrated that the TRAIL-R2 agonistic antibody DS-8273a selectively depleted PMN-MDSCs and M-MDSCs without affecting other immune cell types ^81^.

Our study provides valuable insights into the pathological expansion of MDSCs during HIV-1 infection, utilizing a humanized mouse model. By targeting the TRAIL pathway and depleting MDSCs, we have demonstrated the potential to mitigate HIV-1-mediated immune dysfunction and enhance the function of cytotoxic cells. Our findings not only shed light on the intricate interplay between MDSCs and the immune system during acute HIV-1 infection, but they also offer promising therapeutic avenues for combating HIV-1 infection. The ability to specifically target and deplete MDSCs through the TRAIL pathway represents a novel approach with the potential to improve CD4^+^ T cell reconstitution. Moreover, these findings open doors for further investigations into the utilization of TRAIL-R2 agonistic antibodies in combination with other therapeutic interventions, such as therapeutic vaccines, immune checkpoint blockade, or cell infusions, to enhance the overall efficacy of HIV-1 treatment strategies. Additionally, the insights gained from this study will prompt future exploration of MDSCs and their regulatory mechanisms in other next-generation humanized mouse models as well as SIV-infected macaques, which may further our understanding of MDSC expansion during chronic HIV-1 infection. Collectively, our findings contribute to advancing our knowledge of MDSC biology during HIV-1 infection and provide a foundation for the development of innovative therapeutic approaches that hold promise for improving immune function and ultimately achieving better outcomes in individuals living with HIV-1.

## Materials and Methods

### Sex as a Biological Variable

Humanized mice are generally female, as this generally prevents graft vs. host disease upon humanization of the immune system. However, the CD34+ cord blood derived hematopoietic stem cells (HSCs) are from umbilical cords attached to either male or female newborns. We have not noticed any differences in our results based on the sex of the newborn from whose cord blood unit the HSCs were derived to humanize mice.

#### Statistics

Statistical analysis was performed using GraphPad Prism 9 (GraphPad Software, San Diego, CA, USA). The Student’s t-test was used when comparing two groups, while the two-way analysis of variance (ANOVA) was used when comparing three or more groups.

#### Study approval

The research project indicated above was reviewed and determined not to be “human subjects” research according to the definition provided in 45 CFR 46 102(e)(1) by the Office for the Protection of Research Subjects at The Scripps Research Institute and at The Jackson Laboratory. All protocols involving the use of experimental animals in this study were approved by The Scripps Research Institute and University of Connecticut Health Science Center Institutional Animal Care and Use Committees and were consistent with the National Institutes of Health Guide for the Care and Use of Laboratory Animals.

#### Data Availability

No transcriptome, genome, or spatial sequencing was performed. No new analysis codes were developed.

### Generation of humanized mice and HIV infection

Female NSG-Tg(IL-15) mice (NOD.Cg-*Prkdc^scid^ Il2rg^tm1Wjl^*Tg(IL15)1Sz/SzJ; Stock No: 030890), and NSG mice were purchased from The Jackson Laboratory and used to generate ‘humanized’ (Hu)-NSG-Tg(IL-15) mice featuring a reconstituted human immune system, per our established protocol ^82^ as follows. NSG-Tg(IL-15) mice were sub-lethally irradiated (2 Gy) at 6–12 weeks of age and intravenously injected with 50,000 human CD34+ hematopoietic stem cells (HSCs) enriched from human umbilical cord-blood (StemExpress, San Diego) using the CD34 MicroBead Kit (Miltenyi Biotec, Bergisch Gladbach, Germany). The extent of immune system humanization in each animal was determined by flow cytometric analysis of PBMCs from peripheral blood collected from the submandibular vein twelve weeks after irradiation. PBMCs were isolated by density gradient centrifugation using Ficoll-Paque (GE Healthcare, Chicago, IL, USA), per the manufacturer’s protocol. Cells were washed and resuspended in 0.2 mL PBS supplemented with 2% FBS, incubated with Mouse and Human BD Fc Block Reagent (BD Biosciences, Franklin Lakes, NJ, USA) for 10 min at room temperature (RT), and then stained with antibodies specific to murine CD45 (30-F11; BioLegend, San Diego, CA, USA) and human CD45, (H130; BioLegend), CD3, CD56, and CD19 followed by flow cytometric analysis (see Flow Cytometry description below). Frequencies of human B cells (mCD45^-^hCD45^+^CD19^+^), T cells (mCD45^-^hCD45^+^CD3^+^CD56^-^CD19^-^), and NK cells (mCD45^-^hCD45^+^CD3^-^CD56^+^CD19^-^) were determined. The following criteria were used to confirm robust immune reconstitution: frequency of human CD45^+^ cells ≥ 60% of the total number of human + mouse CD45^+^ cells; and amongst the human CD45^+^ cells, the frequency of B cells is ≥ 10%, T cells is ≥ 15%, NK cells is ≥ 5%, and NK-T cells is ≥ 0.2%. Mice not meeting these criteria were excluded from the experiments. Upon confirmation immune reconstitution, Hu-NSG-Tg(IL-15) mice were infected with 10^5^ infectious units (IU) of HIV-1 strain Q23.17 by intraperitoneal (i.p.) injection. Viral titers were quantified every 7 days by RT-qPCR thereafter until euthanasia.

### Flow Cytometry

Immune enumeration and phenotyping were performed per our established protocol ^82^ as follows. Peripheral blood, spleen, and liver were collected from mice at the designated experimental endpoints. Single-cell suspensions were prepared by mechanically disrupting the tissues through a 40-μm nylon mesh. Immune cells were isolated by density gradient centrifugation with Ficoll-Paque at a density of 1.077 g/mL (GE Healthcare), following the manufacturer’s protocol, and the PMN-MDSC-enriched low-density fraction was retrieved. The immune cells were then washed once in PBS with 2% FBS, resuspended in PBS with 2% FBS, incubated with Murine and Human Fc Block (BD Biosciences) for 10 minutes at room temperature, and stained with antibodies listed in Supplementary Table S1. For FOXP3 staining, cells were permeabilized and stained using the Foxp3/Transcription Factor Staining Buffer Kit (Tonbo Biosciences, San Diego, CA, USA) according to the manufacturer’s guidelines. Cytometric data were acquired using the Cytek™ Aurora flow cytometer (5 L 16UV-16V-14B-10YG-8R model) from Cytek Biosciences (Fremont, CA, USA). The FCS files were analyzed in FlowJo (BD Biosciences), and data visualization and statistical analyses were performed in GraphPad Prism (San Diego, CA, USA).

### Antibody infusions

To deplete human NKp46-domain 1-containing NK cells and CD8a^+^ cells, mice received 100 μg of antibody against human NKp46-domain 1 (clone 9E2; BioLegend) and/or human CD8α (OKT-8; Bio X Cell, Lebanon, NH, USA) by intraperitoneal (i.p.) injection in a volume of 200 µl 3 days before HIV-1 infection and then weekly post-infection for 8 weeks. To deplete human MDSCs, mice received 100 µg of antibody against human TRAIL-R2 (MAB631; Bio-Techne R&D Systems, Minneapolis, MN, USA) by i.p. injection in a volume of 200 µl starting 3 days post-infection and then weekly for 8 weeks. Control mice were injected with vehicle alone.

### Quantitation of HIV-1 viral load by RT-qPCR

Quantitation of HIV-1 viral load by RT-qPCR was performed per our established protocol ^82^ as follows. Mouse peripheral blood was drawn by retro-orbital bleeding into glass capillary tubes. After 30 min at room temperature, 100µL of serum was isolated by centrifugation at 1000 × g for 15 min and viral RNA extracted using the QIAamp MinElute Virus Spin Kit (Qiagen, Hilden, Germany) following the spin column protocol. 50µL of RNA was eluted in Buffer AVE of which 5uL (0.5µg) of RNA was subjected to RT-qPCR in triplicate using qScript XLT 1-Step RT-qPCR ToughMix (Qantabio, Beverly, MA, USA) with the primers/probe 5′-CCTGTACCGTCAGCGTTATT-3′, 5′-CAAAGAGAAGAGTGGTGGAGAG-3′, and 5’-/56-FAM/TGCTTCCTG/ZEN/CTGCTCCTAAGAACC/3IABkFQ/-3′ IDT, Coralville, IA, USA).

## Author Contributions

SP performed funding acquisition. SP and SA performed conceptualization. SA and SP performed formal analysis. SA, KF, and AG conducted the investigation. The methodology was designed by SA and SP. SP performed project administration and supervision. SP provided resources. SA and SP performed validation. SA and SP performed visualization. The writing of the original draft was performed by SA, KF, and SP. All authors contributed to the article and approved the submitted version.

## Acknowledgements

This work was supported by NIH grants R01AI161014 and R21AI170555 to SP and seed funds to SP from The Scripps Research Institute and the Jackson Laboratory for Genomic Medicine.

## Supplemental Materials

**Supplementary Table 1:** Flow Cytometry Antibodies. * Murine specific antibodies, all others are specific to human antigen

| Target | Fluorophore | Clone | Catalog Number | Company |
| --- | --- | --- | --- | --- |
| CD45 | BV605 | HI30 | 304042 | Biolegend |
| CD45 | BUV395 | HI30 | 563792 | BD |
| CD56 | BV570 | HCD56 | 318330 | Biolegend |
| CD16 | PE-Cy5 | 3G8 | 302010 | Biolegend |
| CD3 | APC/Fire 750 | UCHT1 | 300470 | Biolegend |
| CD8a | BUV395 | RPA-T8 | 563795 | BD |
| CD8a | BUV563 | RPA-T8 | 612914 | BD |
| CD8a | BV785 | RPA-T8 | 301046 | Biolegend |
| CD19 | BV750 | HIB19 | 302262 | Biolegend |
| CD11b | PE/Dazzle 594 | ICRF44 | 301348 | Biolegend |
| CD14 | BV510 | MΦP9 | 563079 | BD |
| CD66b | PE | G10F5 | 305106 | BD |
| Tbet | BV711 | 4B10 | 644820 | Biolegend |
| EOMES | PerCP-eFluor 710 | WD1928 | 46-4877-42 | Invitrogen |
| TNF-a | BV750 | MAb11 | 566359 | BD |
| IFN-g | AF700 | 4S.B3 | 502520 | Biolegend |
| Perforin | PE | B-D48 | 353304 | Biolegend |
| Granzyme B | PacBlue | GB11 | 515408 | Biolegend |
| CXCR6 | PE/Dazzle 594 | K041E5 | 356016 | Biolegend |
| CD62L | PerCP-Cy5.5 | DREG-56 | 304824 | Biolegend |
| CD57 | BB515 | NK-1 | 565285 | BD |
| CD69 | BV510 | FN50 | 310936 | Biolegend |
| CD107a/Lamp1 | BV786 | H4A3 | 563869 | BD |
| CD4 | BV650 | OKT4 | 317436 | Biolegend |
| CD4 | BV750 | SK3 | 344644 | Biolegend |
| CD8a | BV785 | RPA-T8 | 301046 | Biolegend |
| CD11b | PE/Dazzle | ICRF44 | 301348 | Biolegend |
| CD14 | BV510 | 63D3 | 367124 | Biolegend |
| CD15 | Pacific Blue | W6D3 | 323022 | Biolegend |
| CD16 | PE/Cy7 | 3G8 | 302010 | Biolegend |
| CD33 | FITC | HIM3-4 | 303304 | Biolegend |
| CD66b | PE | G10F5 | 305106 | Biolegend |
| CD163 | BV711 | GHI/61 | 333630 | Biolegend |
| PD-1 | BV480 | EH12.1 | 566112 | BD Biosciences |
| NKG2D | Super Bright 436 | 1D11 | 62-5878-42 | ThermoFisher |
| FOXP3 | Alexa Fluor 532 | PCH101 | 58-4776-42 | ThermoFisher |
| HLA-DR | BV650 | L243 | 307650 | Biolegend |
| FoxP3 | AF532 | 2H7 | 58-0209-41 | Invitrogen |
| mCD45* | AF700 | 30-F11 | 103128 | Biolegend |
| mCD45* | PerCP/Cy5.5 | 30-F11 | 103132 | Biolegend |
| Human Fc Block <sup>TM</sup> | N/A | N/A | 564220 | BD Biosciences |
| Mouse Fc Block <sup>TM</sup> | N/A | 2.4G2 | 553142 | BD Biosciences |

**Supplemental Figure 1:**
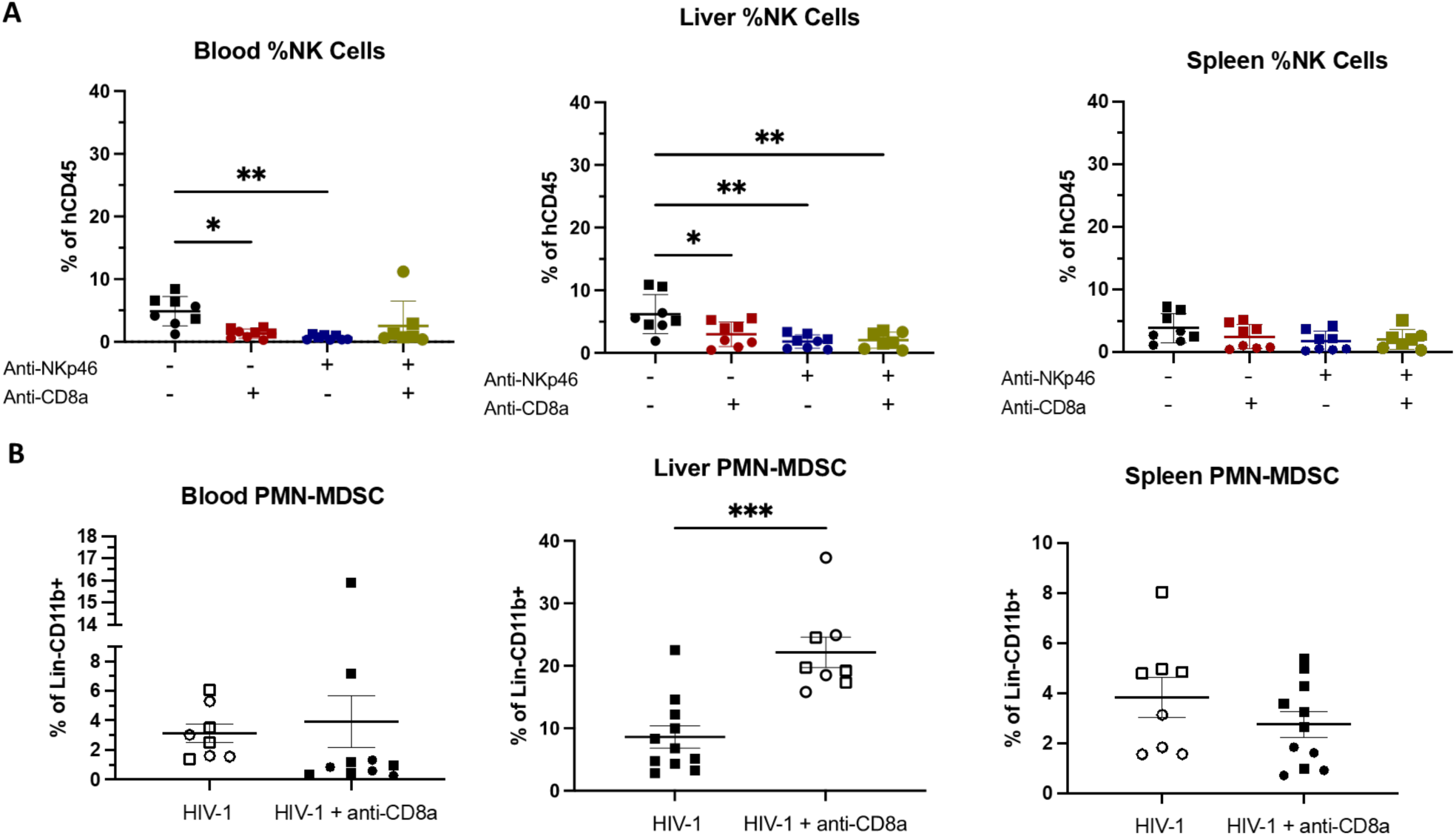
A) Percentage of NK cells (hCD45+mCD45-CD3-CD56+) as a percentage of human immune cells (hCD45+mCD45-) in HIV-1 infected mice intraperitoneally injected with either Anti-CD8a+ (OKT8), Anti-Nkp46 (9E2), both or neither antibody mice (2 donors, n=3-4 per experimental group). B) Frequency of PMN-MDSC in blood, spleen, and liver of HIV-1 infected mice with or without depletion of CD8a+ by weekly intraperitoneal injection of Anti-CD8a (OKT8). Statistical significance was calculated using an unpaired t-test or ANOVA with multiple comparisons *P < 0.05, **P < 0.01, **P < 0.005, and ****P < 0.0001.

**Supplemental Figure 2:**
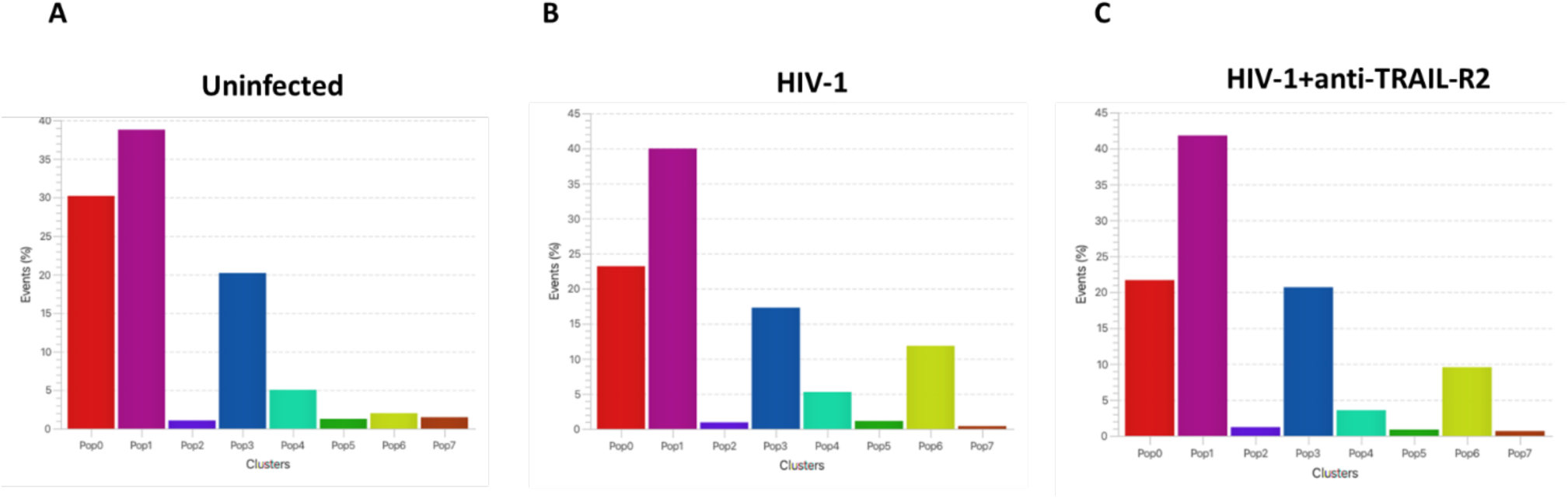
Percentage of cells in each cluster generated from unsupervised FlowSOM clustering. A-C) t-SNE dimensionality reduction of hCD45+mCD45-cells positive for either CD3 and/or CD56 with overlayed with FlowSOM clusters of splenic immune cells Hu-NSG-Tg(IL-15) mice 8-weeks post HIV-1 infection, with or without treatment with anti-TRAIL-R2 agonistic antibodies and uninfected donor-matched control mice. Cells from individual mice were pooled and downsampled to 50,000 events prior to t-SNE and FlowSOM analysis.

## Notes

Conflict of Interest Statement: The authors have declared that no conflict of interest exists.

### Competing Interest Statement

The authors have declared no competing interest.

